# Holmes-ITS2: Consolidated ITS2 resources and search engines for plant DNA-based marker analyses

**DOI:** 10.1101/263541

**Authors:** Hongjun Li, Hong Bai, Shaojun Yu, Maozhen Han, Kang Ning

**Author notes:** These authors contributed equally to this work (Hongjun Li, Hong Bai).

## Abstract

Plants are valuable resources for a variety of products in modern societies. Plant species identification is an integral part of research and practical application on plants. In parallel with high-throughput sequencing technology, the high-throughput screening of species is in high demand. Highly accurate and efficient DNA-based marker identification is essential for the effective analysis of plant species or biological constituents of a mixture of plants as well. Therefore, it is of general interests and significance to generate a comprehensive and accurate DNA-based marker sequence resource, as well as to build efficient sequence search engines, for the accurate and fast identification of plant species.

In this work, we have firstly established a high-quality ITS2 sequence database of plant species containing more than 150,000 entries, through the systematical collection and manually collation of the published ITS2 sequencing data of plant species, data quality control, as well as representative sequence refinement based on clustering method. Secondly, an accurate and efficient plant species identification system based on ITS2 sequence was constructed, which is the proper combination of sequence search algorithms including BLAST and Kraken. Through the deployment of high-performance and frequently updated web service, it’s expected to serve for a wide range of researchers involving the taxonomy classification of plant species, as well as for deciphering of plant mixed systems including herbal materials in TCM preparations.

The Holmes-ITS2 web service is freely accessible at: http://its2.tcm.microbioinformatics.org/. The input of this web service could be multiple sequences in a single fasta format, to search for matching ITS2 biomarker sequences already annotated in the database. This sequence-based search is based on two engines: BLAST, and k-mer based Kraken. Alternatively, users can directly search for species name for the corresponding ITS2 biomarker sequences. The web service has been put to the test by more than 50 experts from China, Denmark and US, and the average running time for the search ranges from 3-30 seconds for up to 100 sequences as a batch query.

## INTRODUCTION

Currently, there are more than 300,000 plant species on earth that have been described (1), providing valuable resources such as food, fibers, timber and medicine, etc. to support modern societies (2). Plant species identification and taxonomy classification are the basis of ecology, botany and biology, especially related with utilization and protection of plant resources. On one hand, for those plants for consumption, including various food and drugs, the accurate identification of plants is requisite to avoid the safety issues caused by the misuse of closely related species or adulteration and ensure the theoretical efficacy (3). Researches on plants would be carried out with the aid to the knowledge accumulated through the deep examination of the plants. On the other hand, as for researches on biodiversity and conservation of endangered fauna and flora, building accurate knowledge-base of plants is essential for their rational protection, which would also aid for preventing illegal trade of endangered plants (4). Therefore, accurate and rapid identification of plants would be essential for safe and rational utilization of plant resources and effective study and protection of plant biodiversity.

Besides traditional approaches to identify plants through physical characteristics or inference from chromatographic fingerprints generated by High Performance Liquid Chromatography (HPLC) or Thin Layer Chromatography (TLC) technologies, which bring difficulties to differentiate species with indistinguishable or changed morphology and chemical constitutions. DNA-based molecular markers were introduced to be an efficient and reliable means of identification of plant species(5), especially in mixtures that contain more than one species(6). DNA barcodes are based on a standardized short sequence of DNA from a small region of a species’ genome that can distinguish the species from others in the same kingdom quickly and accurately(7). As a representative marker, the internal transcribed spacer 2 (ITS2) is a fast-evolving locus of the nuclear rRNA cistron which has large variations in sequences also with features as easy amplification and high universality and is thus appropriate to be a proper DNA barcode for studies and inferences of phylogenies at low taxonomic levels(8). For plants, ITS2 has been broadly used as an effective DNA-based marker for identification of organisms at species or sub-species level(9). As the development of next generation sequencing (NGS) technology, generation of DNA-based marker sequence data is becoming easier and easier, and the amount of relative data keeps growing, through which, high-throughput research on plant preparations has become a trend.

Compared with a single plant, the identification of mixtures that contain more than one plant species (plant mixed system) is more complicated and challenging. Such identification has practical value for quality evaluation of products in the market made of plant materials, and one typical example of these is the Traditional Chinese Medicine (TCM), which usually contains multiple plant species. The existence of DNA belonging to different plant raw materials makes it possible and convenient to identify plant species in a mixed system through methodologies that could take advantage of DNA-based markers. Identification of a plant mixed system is to recognize the taxonomy species belonging to various raw materials in essence, depending on accurate identification of DNA-based markers of plants including ITS2. Based on high-throughput sequencing and big data mining techniques, metagenomic methods have become one of the most important and effective approaches to understand and analyses the structure and functionality of a biological mixture(10), which could help to establish an accurate and efficient method for biological constituent analysis of the plant preparation or the plant mixed system.

The requirements of high-throughput analyses of biological constituents of plant preparations or plant mixed systems put forward a very high standard for the precision (precision to species or subspecies level), accuracy (low false-positive rate) and efficiency (processing batch and bulk data quickly) of identification and comprehensive species coverage. The existing databases of species identification of plants such as TCMBarcode(11) and ITS2 Ribosome RNA Database(12) are more focused on the analysis of single sequence data in respect of identification of DNA-based marker sequences, which is not adapted to researches with high throughput sequencing data of plant DNA-based marker. By using ITS2 as DNA-based marker and with the help of metagenomic methodologies, we designed and constructed a plant DNA-based marker database and taxonomy classification and organism identification system, Holmes-ITS2, to serve for high standards for the identification of biological constituents of plants (website: http://its2.tcm.microbioinformatics.org/). Through the process of raw data collected and the optimization of search algorithms, accurate and efficient identification of plant species could be achieved to match the high throughput sequencing data of plant orplant mixture system for research or practical purpose.

## MATERIALS AND METHODS

### Data source and quality filter of raw sequences

Raw ITS2 sequencing data were extracted from the NCBI nucleotide database (https://www.ncbi.nlm.nih.gov/nucleotide/) in Genbank format searched with key words “ITS2”, together with “Species” filtered to “Plants” in April 2016. First, extractions of target ITS2 sequences of data downloaded were carried out with in-house scripts to trim boundaries of each ITS2 sequence, namely 5.8S and 28S rRNA genes which were highly conserved among different plant species (**Figure 1A**). Then, manual picking of sequences was performed to collect positive entries that the script didn’t cover. Due to the absence of ITS2 location annotations of some raw data, these sequences were moved to the candidate dataset first, and then a Hidden Markov Model was trained based on well-annotated ITS2 sequences to predict the potential ITS2 regions of these candidate sequences, before these ITS2 sequences could be included into our curated database. For all ITS2 sequences extracted based on the annotations, quality filter was performed in accordance with criteria as follow (**Figure 1A**): (1) length below 100 bp, (2) length above 900 bp, (3) belonging to reduplicate entries, (4) with more than three ambiguous base pairs, (5) belonging to environment samples or unclassified samples. The quality control steps filtered ITS2 entries with either low sequence quality or obscure taxonomy annotation. Also, as there may be retrieval results with key words while containing no target sequence, which should be screened out.

**Figure 1.**
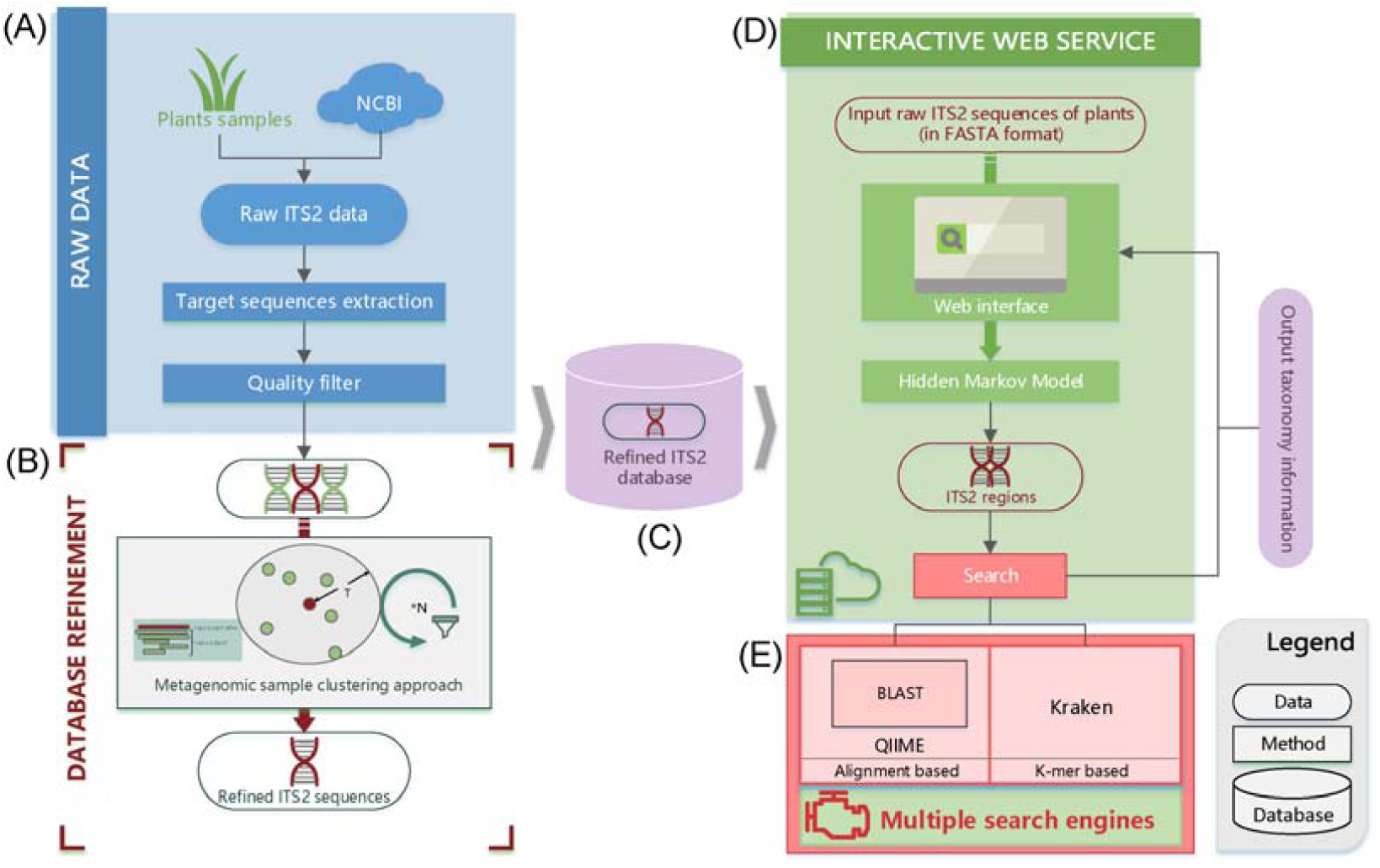
The whole workflow of the Holmes-ITS2 system including database, web service and search engines. **(A)** Raw ITS2 sequences obtained from NCBI, target sequence extraction and sequence quality filtration. **(B)** Database refinement by metagenomic sample clustering and representative ITS2 sequence selection. **(C)** The refined ITS2 database. **(D)** Interactive web service with **(E)** multiple search engines enabled.

### Building and applying Hidden Markov Model

Due to the restriction of primers during amplification processes, ITS2 sequences obtained from the actual experiments usually contain sequences out of both boundaries, namely 5.8S and 28S rRNA genes in eukaryotes. With 1,000 sequences representing clean and complete ITS2 without ambiguous base pairs and out-of-boundary sequences from the data set passing the quality filter, a Hidden Markov Model of ITS2 sequences was trained through multiple sequence alignments with MUSCLE(13) (Version 3.8.31) and HMMER3(14) (Version 3.1b2) to build the model with default parameters. The model was then applied to predict the boundaries of ITS2 regions(15) of the candidate data set to extract target ITS2 sequences through the HMMER3 program. Through the search process based on probabilistic inference, those potential ITS2 sequences in candidate data set could be extracted. These predicted sequences were also filtered accordance with the criteria as set above. For sequences itself or whose sub-sequences that do not match the model, they were considered as non-ITS2 sequences and filtered out.

### Metagenomic sample clustering approach to refine the ITS2 sequence database

Raw nucleotide sequences in NCBI might have a problem of their identity: taxonomy information recorded of some sequences was not accurate or differed greatly of some sequences with high similarity. These problems would lead to deviation of organismal identification and taxonomic classification based on sequence similarity. To realize fault-tolerance and reduce impact of the problems caused by original data, as well as to refine the database, sequence clustering approach used in metagenomic sample analysis (namely the UCLUST algorithm) was introduced (**Figure 1B**), which was generally used in a different context to generate clusters of (uncultivable or unknown) microorganisms (Operational Taxonomical Unit, OTU), grouped by DNA sequence similarity of a specific taxonomic marker gene(16). In the clustering process, sequences whose similarities above a certain value were grouped into a cluster expected to belong to the same species or closely related species.

The sequence cluster procedure was carried out by the UCLUST program (version v1.2.22q). Sequences were sorted through their length first and processed in order one by one. If a sequence being processed matched an existing centroid, it was assigned to that cluster, otherwise it became the centroid of a new cluster(17). The similarity threshold of the ITS2 sequence was set to 0.99, so that highly homologous plant species could be clustered.

After the clusters were generated each containing highly similar sequence, we have performed further filtration for each cluster. Sequences whose phylogenetic relationships of species annotated diverging obviously in a cluster would be processed by the principle that isolated sequences (below 10% of the total sequences in the cluster) would be filtered while a dominant species in a majority number (above 90%) of the total sequences in the cluster would be retained. Finally, sequences with inaccurate annotations were filtered out.

### Deployment and parameter setting of multiple search engines

As the taxonomic classification of applying the database was based on sequence similarity search, for the consideration of high accuracy and efficiency, multiple search engines were designed as **Figure 1E**. For alignment-based BLAST(18) working in QIIME(19), data was formatted as two separated files containing sequences and taxonomy information, respectively. The mapping relation of ITS2 sequences and their species information was assigned. An efficient algorithm of sequence search, k-mer based Kraken(20), a fast and accurate algorithm initially used for assigning taxonomic labels to metagenomic DNA sequences, was also applied as an efficient species classification method. The core of Kraken was a database containing records consisting of a k-mer and the LCA (the least common ancestor) of all organisms whose genomes contain that k-mer(20). Sequences were classified by querying the database for each k-mer in a sequence, and then using the resulting set of LCA taxa to determine an appropriate label for the sequence. As for Kraken, data was formatted (aligned to generate k-mers contained within the database used for Kraken) with built-in commands, and the NCBI taxonomy database was adopted as taxon information (mapped to k-mers with the GI number) for the construction of the Kraken custom database.

### Database Comparison

With the existing ITS2 databases (**Table 1**) as reference, the performance of Holmes-ITS2 database was tested from three aspects, including accuracy, efficiency and data coverage. For accuracy test, considering that the existing two databases couldn’t support submission of batch data (specifically, one sequence once submission of TCMBarcode and five of ITS2 Ribosomal RNA Database), only a small number (1,000 entries) of raw ITS2 sequences was picked at random from NCBI for testing a purpose. To test and compare the accuracy rate of the three databases, the whole dataset was inputted to the taxonomic classification pipeline of Holmes-ITS2 (**Figure 1D**) to get the classification result, and for the online databases, test dataset was submitted manually by the maximum amount of data acceptable to the database batch after batch and results were recorded.

**Table 1.**
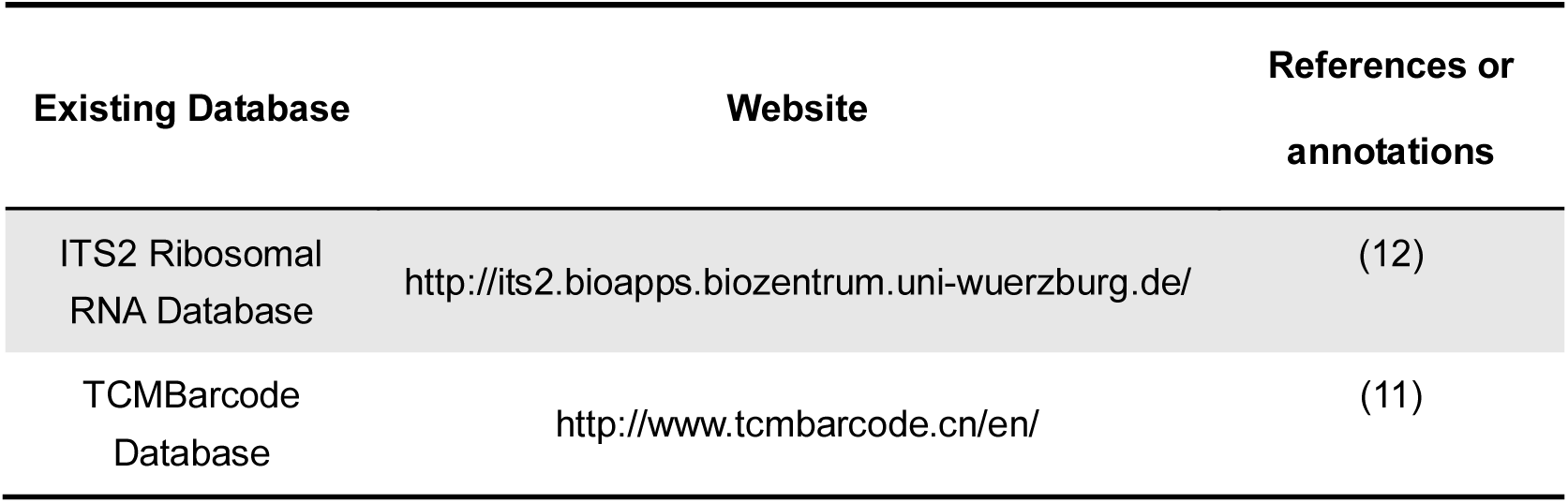
Existing databases selected for database comparison.

For efficiency test, different amount of raw ITS2 queries (from 100 to 200,000 queries) as test datasets were obtained from NCBI. Time cost of two main sections of the identification process, including the HMM (Hidden Markov Model) prediction and classification of target sequences by different search engines was recorded under different data size to make the horizontal and vertical comparison and evaluate the overall identification speed of different databases.

For data coverage of databases, to gather statistics, the number of entries was counted directly for the Holmes-ITS2, and data was collected from the ITS2 Ribosomal RNA website and the TCMBarcode website (**Table 1**), respectively.

### Case study in TCM research

As a typical plant mixed system in practical application, TCM preparations contain medicinal plants as the main raw materials. While misidentification of closely related species, erroneous substitution with other herbs or intentional adulteration would reduce the efficacy or harm to human beings, which were serious security issues. Thus, accurate identification of medical materials was an essential step to reduce or avoid the consequences of such problems. Accurate analysis of biological components of TCM preparations depended on high accurate identification of DNA-based markers. To measure the identification performance of actual TCM preparations of Holmes-ITS2, a case study was carried out based on the sequencing data of our previous research (21), which contributed to analysis the biological ingredients of a classical TCM preparation Liuwei Dihuang Wan (LDW).

The identification was carried on the 454 sequencing data included ITS2 sequences of 3 biological replicates of a reference (RE) and 9 specimens from 3 manufacturers (MH, MS and MT) each with 3 batches (A, B and C). The quality control was performed as the previous study using Mothur(22) (version 1.39.5). Sequences whose length below 150 bp and average quality score below 20 in each 5 bp-window rolling along the whole read were discarded. Those sequences containing uncorrectable barcodes, primer mismatches, ambiguous bases and homopolymer runs more than 8 bases were also removed from the datasets. With reads sorted by tag sequences, they were then identified in accordance with the pipeline of Holmes-ITS2 and summary and statistics on the results were made finally to compare the biological components of LDW specimens commercially available. In order to ensure the consistency of the identification criteria with the previous study, ITS2 sequences for which the corresponding possible species was evidenced by 3 or less reads were filtered and BLAST search was performed with the E-value threshold set to 1E-10.

### Construction of web service

In order to facilitate the utilization of the Holmes-ITS2 species identification services, the web service of Holmes-ITS2 taxonomy identification system was designed and built (**Figure 1D**) based on a high-performance computing platform. The web service features including species ITS2 sequence browsing and multi-search engines for homology search of existing sequences (**Figure 1E**).

The underlying infrastructure of the web service was based on the typical LNMP architecture and the PHP framework Laravel. The running mechanism (**Figure 1D**) was that as a batch of fasta sequences of a TCM preparation sample was pasted as input (data in a submission to the web server was regarded as a sample), HMM model was applied to predict the ITS2 region of each sequence (Originally, it was supposed that the ITS2 region was a part on the sequence to be queried). Target ITS2 sequences would be extracted from raw queries before they will be passed to the search engine selected. For each sequence failed to be predicted, the raw sequence would also be passed to the search engine, and it would try to identify the taxonomy of the sequence. After the identifications of all sequences were completed, an analysis report would be shown including taxonomy classified of each sequence and the statistics results of species constituents of the sample (including genus-level and species-level results).

### Backend data update mechanism

As the development of high-throughput sequencing technology, nucleotide data of species which TCM preparations comprise was exploding. In order to maintain maximum data coverage, to be specific, the plant species coverage, the data update mechanism was designed. For a defined period (usually every six months), new ITS2 sequences released on the NCBI nucleotide database would be extracted (entries sorted by ‘Data Released’) and processed in accordance with the methods designed and standards set as explained previously. Consistency of newly released data would be ensured through the metagenomic sample clustering approach with all data that was already in the database. Finally, new entries meeting the standard would be appended into Holmes-ITS2.

With updates performed periodically, newly released raw ITS2 sequencing data could be possessed and be included into Holme-ITS2 to maximize data coverage and database availability.

## RESULTS

### Quality filtration improves the sequence quality

The current dataset for plant DNA-based marker based on ITS2 consisted of 169,950 sequences in genbank format from the NCBI nucleotide database in April 2016. With the annotation information and the predictions by the HMM model. Out of these sequences, 162,544 sequences’ (96.78% of raw data) ITS2 region was extracted (**Figure 2A** and **Figure 2C**). After all quality filtration steps, 986 sequences (0.58%) with more than three ambiguous base pairs, 953 reduplicate sequences (0.56%), 565 sequences (0.33%) with bad annotation, including “environmental samples” or “unclassified”, and 1190 sequences (0.7%) with length above 900 bp or below 100 bp (accordance with **Figure 2B**, 99.26% of the total that included complete or partial sequences distributing in the limited region) were screened. In terms of the length of ITS2 sequences, which mainly distributed throughout the region from 200 base pairs to 300 base pairs with the peak located at the sequence length of 221 base pair, the interval cut-off was set from 100 base pairs to 900 base pairs, expected for full-length ITS2 and better performance of the cluster analysis. Overall, the maximum of data loss occurred in the trimming procedure of candidate entries with unclear annotation of gene position carried out manually (2.50%) while the minimum in the filter of entries with imprecise taxonomy annotation (0.35%), as each percentage counted was based on the result of the previous procedure. Finally, 158,850 sequences (93.5%) were retained for cluster analysis.

**Figure 2.**
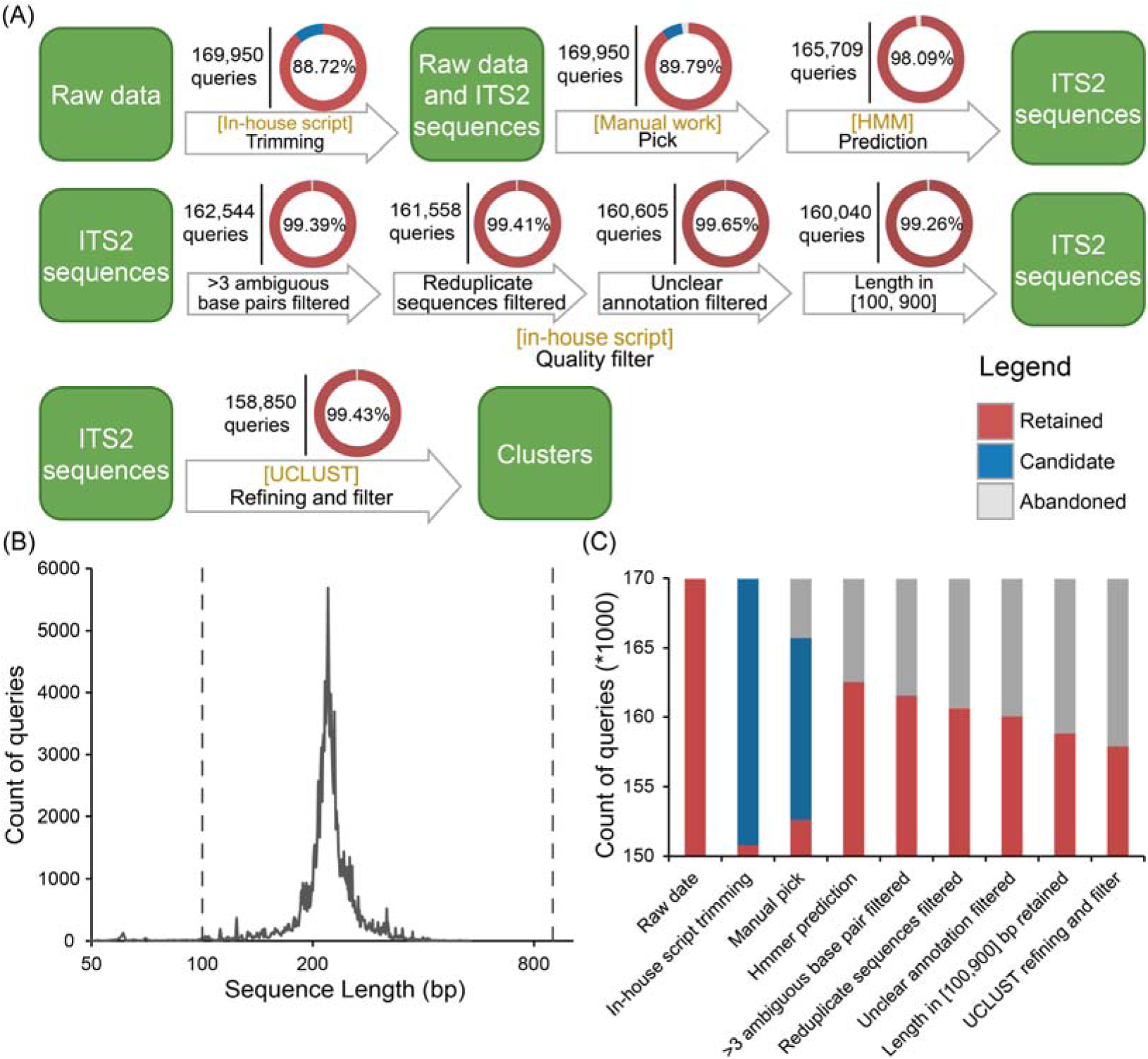
Statistics results of for each step in the procedure of data processing and sequence length distribution. **(A)** The number located on the left of the vertical line above the arrow represents the initial number of queries in each processing step. Pie chart displays the composition and change of the dataset during each sub-step of processing. Red fan representing the percent of retained queries, blue fan of candidate queries which would be treated in the next step (which might be filtered) and blank fan of queries filtered within the step. **(B)** The statistics result of length distribution of ITS2 sequences extracted and the filter thresholds, between which there were more than 90% of total sequences. The length of abscissa axis was displayed as the logarithm of the actual length to base 4 **(C)** Stacked column chart displaying the number of entries and its trend over the processing procedures.

### Hidden Markov Model further improves the quality of entries

1,000 high-quality sequences were selected from the data set after the quality filter to perform multiple alignments with MUSCLE program. These sequences would meet the following requirements: (a) complete ITS2 sequence with length in range from 200 bp to 300 bp, (b) without ambiguous base pairs, and (c) with clear taxonomy annotation,. An HMM model was constructed for ITS2 from the alignment result with HMMER. The candidate dataset after manual pick, which has to lack of the ITS2 gene annotation, was predicted by the HMM model to extract potential ITS2 sequences for maximum utilization of the data. By utilizing the trained model on the 13,116 candidate queries, 75.87% (9,951) potential ITS2 sequences were predicted based on the HMM model (**Figures 2A** and **2C**), which has actually improved the availability of the overall dataset.

### Metagenomic sample clustering approach refines the sequence set

According to the principle that sequences with high similarity of the same gene belong to the same or closely related species, a metagenomic clustering analysis based on sequence similarity was carried out on the dataset after quality filter by UCLUST program (**Figure 1B**). With the threshold of sequence similarity set to 0.99 (sequences with similarity above 0.99 would be assigned to the same cluster), 36,765 clusters were aggregated. In theory, sequences within the same cluster should belong to the same or closely related species (belonging to the same genus). By careful manual check, 913 sequences with taxonomy annotated not correspond to the taxonomy represented by most sequences (over 90%) in their clusters were screened out (**Figures 2A and 2C**), which may have influence on the classification. Finally, sequences that were left were highly consistent within each cluster, and a refined and highly consistent dataset was obtained.

After the processing and filtering procedures, there were 157,937 clean plant ITS2 sequences (92.93% of raw data) with high-quality bases and taxonomy annotations. The taxonomy hierarchical structure to the database was shown as **Figure 3**, in which there were 2 phyla, 38 classes, 169 orders, 501 families, 8,385 genera and 65,281 species, uniquely.

**Figure 3.**
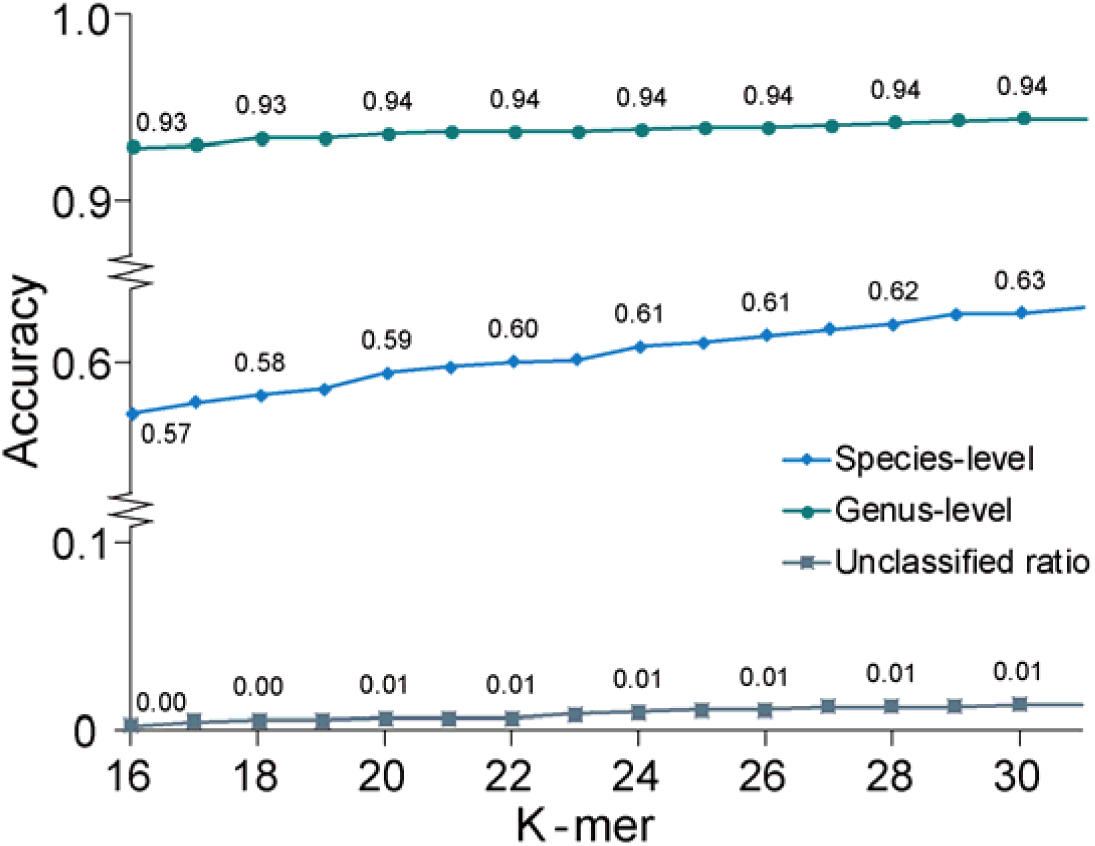
Classification accuracy comparison based on different k-mers for Kraken search engine. As the increase of k-mer set for the database, the accuracy of Kraken search has shown a growing trend at different levels, with a slightly increase for unclassified results, which was the basis for choosing the final k-mer value.

### Multiple search engines improve search accuracies

Two search engines were deployed for organismal identification and taxonomic classification with the database (**Figure 1E**). For the BLAST working in QIIME, an in-house script was used to format the data in order to enable the software to invoke directly. For the Kraken custom database, NCBI taxonomy containing the GI number to taxon map, as well as the taxonomic name and tree information was downloaded in September 2016. The database was built by the built-in command of Kraken. And it was noted that when the k-mer was adjusted to 31-mer it showed the relatively best performance with the proportion of sequences unclassified rising slightly (**Figure 4**). As a result, for the best classification accuracy, the k-mer in the Kraken custom database was adjusted to 31-mer.

**Figure 4.**
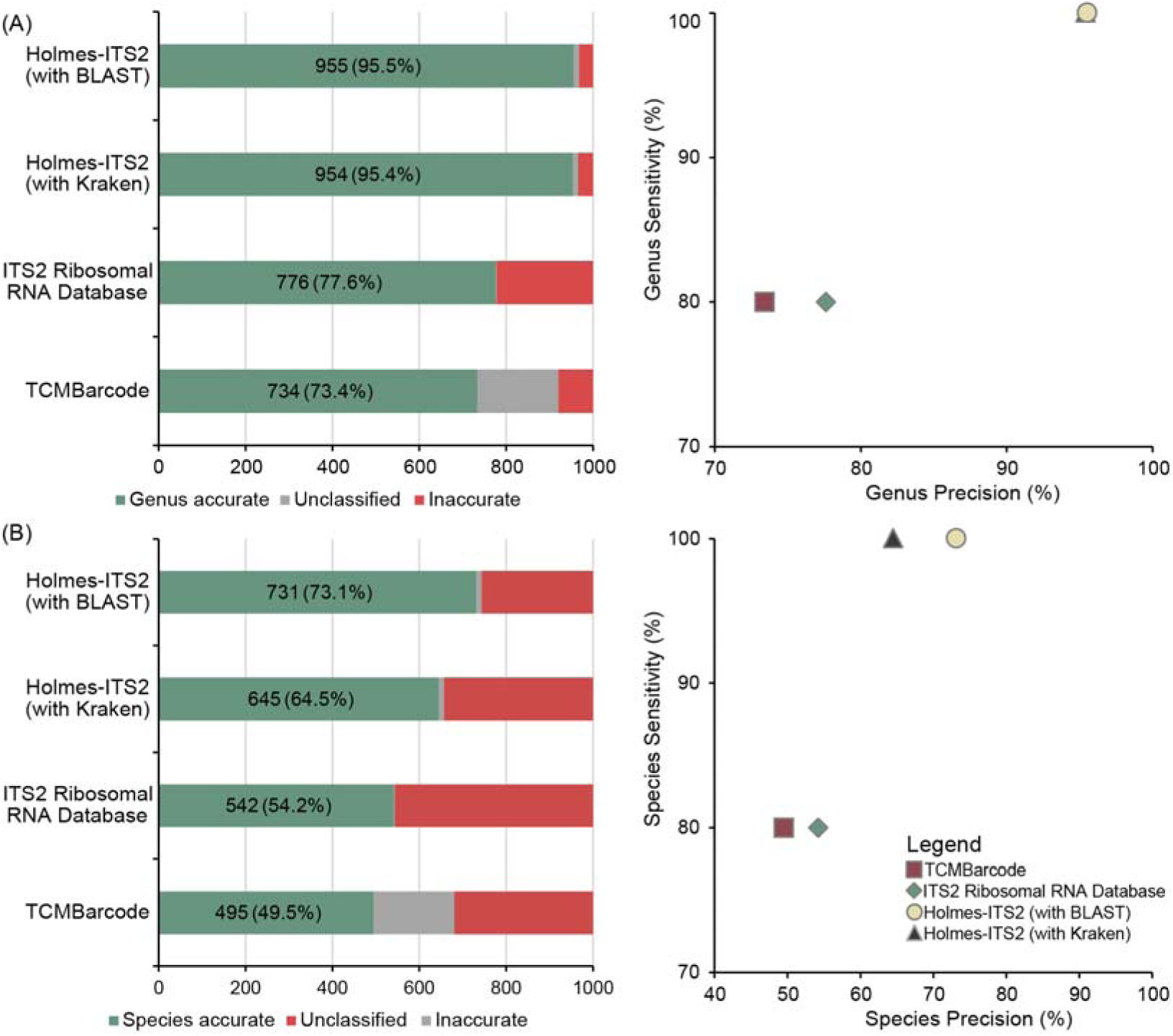
Comparison of classification accuracy, precision and sensitivity of three ITS2 databases. **(A)** Genus-level accuracy, precision and sensitivity and **(B)** species-level accuracy, precision and sensitivity were shown for three databases. For each of the horizontal histogram, the number within each bar representing the number and rate of sequences classified correctly in target taxonomy level. For each of the scatter plot, x-axis represents the species identification precision, and y-axis represents the species identification sensitivity.

### ITS2 database and search system comparison

#### Classification accuracy

To compare Holmes-ITS2’s accuracy to the other two databases (**Table 1**), 1,000 raw entries as testing dataset were selected randomly from NCBI was classified and genus-level and species-level accuracies, sensitivity and precision were measured (**Figure 4**). Here, as Holmes-ITS2 would only return the best-match taxonomy of a sequence while there may be more than one result scoring equally of the two databases. It was considered a right classification only if the real taxonomy of a sequence was in the best-match result set of the classification. Furthermore, sensitivity referred to the ratio of queries assigned to the correct genus or species, and precision referred to the ratio of correct classifications in genus-level or species-level out of the total number of classifications tried.

The classification accuracy, precision and sensitivity of three databases were investigated in genus-level and species-level, respectively. For genus-level accuracy and precision, Holmes-ITS2 database with BLAST as search engine appeared to be the highest of all three databases (**Figure 4A**), which was up to 95.5%. Furthermore, those of Holmes-ITS2 with Kraken as search engines were very close to BLAST, which was within 0.1 percent while TCMBarcode and ITS2 Ribosomal RNA database only did so for 73.4% and 77.6%. For species-level accuracy and precision, all three databases were not that high of the genus-level, which was caused by the resolving power of ITS2 itself. Similarly, Holmes-ITS2 database with BLAST as search engine got the highest classification accuracy and precision (73.1%) among all three databases with a clear superiority. To be specific, there were 8.6%, 18.9% and 23.6% gap between that of Holmes-ITS2 with Kraken as search engine, ITS2 Ribosomal RNA database and TCMBarcode database, respectively. In view of the overall situation, Holmes-ITS2 showed the best performance in classification accuracy among the three databases with certain advantages.

#### Classification speed

As the increase as the data sizes of recent researches, efficiency classification in acceptable time was also an important issue to consider. Time cost to the classification process under a different number of queries was calculated with timing datasets including the different amount of sequences created with raw queries obtained from NCBI. To reduce the accidental error, the time cost was measured by averaging the results of tests under a certain number of sequences repeated 5 times and carried out on the server during off-hours. For the other two ITS2 databases, since the online service supported only a small number of sequences submitted for classification each time, the classification speed test was only carried out on Holmes-ITS2 in the period of HMM prediction and taxonomy classification with different search engines. This was also a short board in existing ITS2 databases, which was not appropriate for actual applications aiming at high-throughput data.

Classification speed of two main procedures was evaluated as **Figure 5.** Basically, time cost of HMM prediction (**Figure 5A**) and search by BLAST (**Figure 5B**) showed a linear increasing trend as the increase with the number of queries, and averagely, the speeds of the two procedures were 250 queries and 15 queries per second on our web server. It was noteworthy that the time cost of classification by Kraken, which didn’t rise significantly in the range from the number of queries tested with an average classification speed of more than 6,000 sequences per second. During classification of the small amount of data, the fluctuation of time cost was caused by that it was far from the maximized classification speed of Kraken.

**Figure 5.**
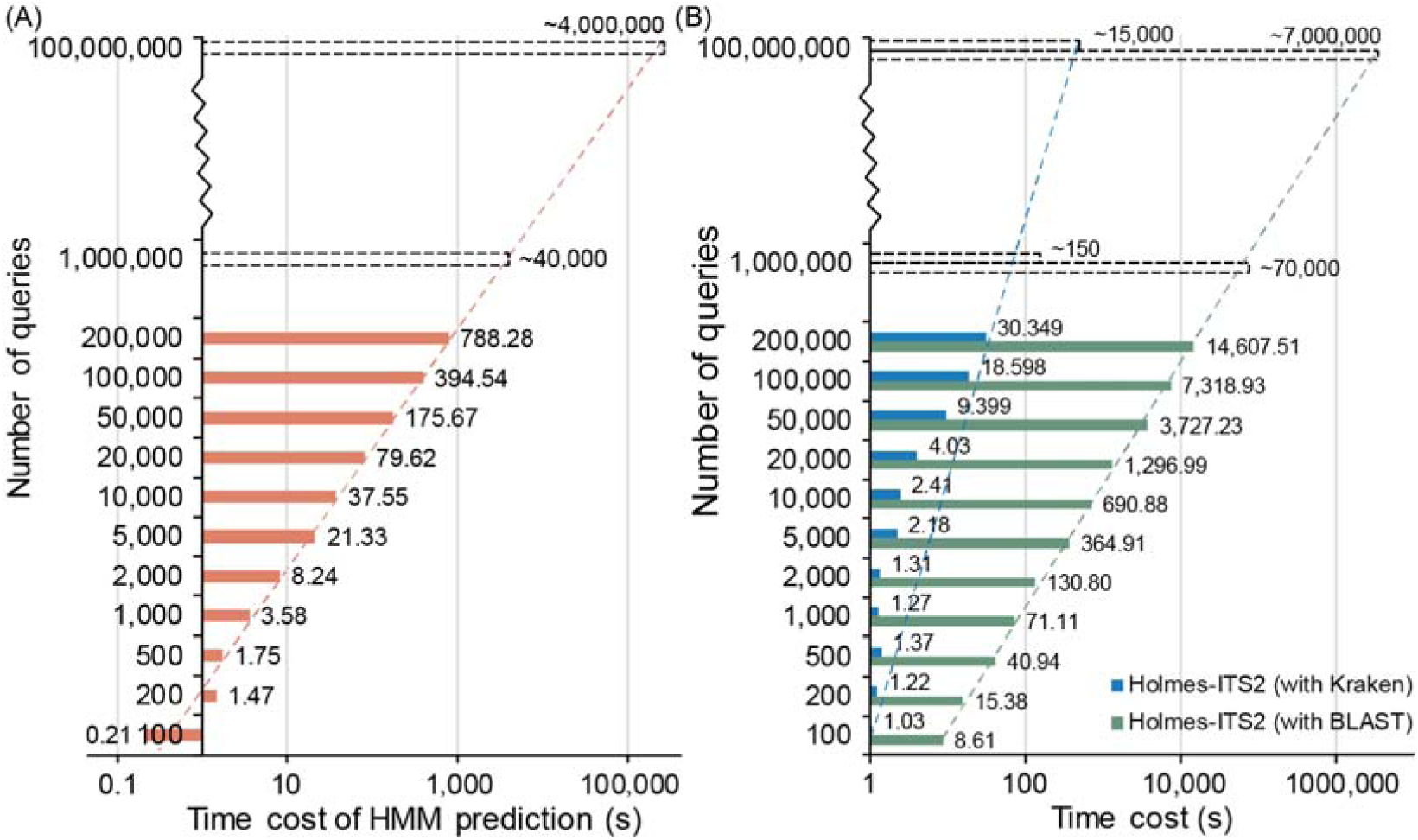
Classification speed comparison of Holmes-ITS2 with BLAST and Kraken as search engines on the test datasets with different number of queries. Two most time-consuming partsto the classification process, including **(A)** HMM prediction and **(B)** identification with BLAST and Kraken search engines. Each solid horizontal bar represented the time cost in a procedure of the whole classification under a different number of queries. Trend lines of time cost of different procedures were shown by broken lines. The classification time cost of different step remained a linear increase trend in general and thus the time cost under 1 million and 1 hundred million queries were estimated shown as dotted bars.

In general, classification with Kraken had a significant advantage over the legacy BLAST, especially for the large size of data. Based on the previous result, that was at the expense of a little of classification accuracy. In conclusion, classification with BLAST could yield a more accurate result of taxonomy identification, while the use of Kraken would be able to reduce time consumed greatly by the overall classification procedure, especially for large quantities of sequence data.

#### Analysis of data coverage of databases

As the statistics results shown as **Figure 6** There were 2 phyla, 38 classes, 169 orders, 501 families, 8,385 genera and 65,281 species uniquely in Holmes-ITS2. Compare with ITS2 Ribosomal RNA database (114,733 queries in total belonging to not only plant species but also animals, etc.) and TCMBarcode (12,221 queries for not only plant species, both statistics results were obtained in September 2016). Holmes-ITS2 also had advantages in the overall data coverage of ITS2.

**Figure 6.**
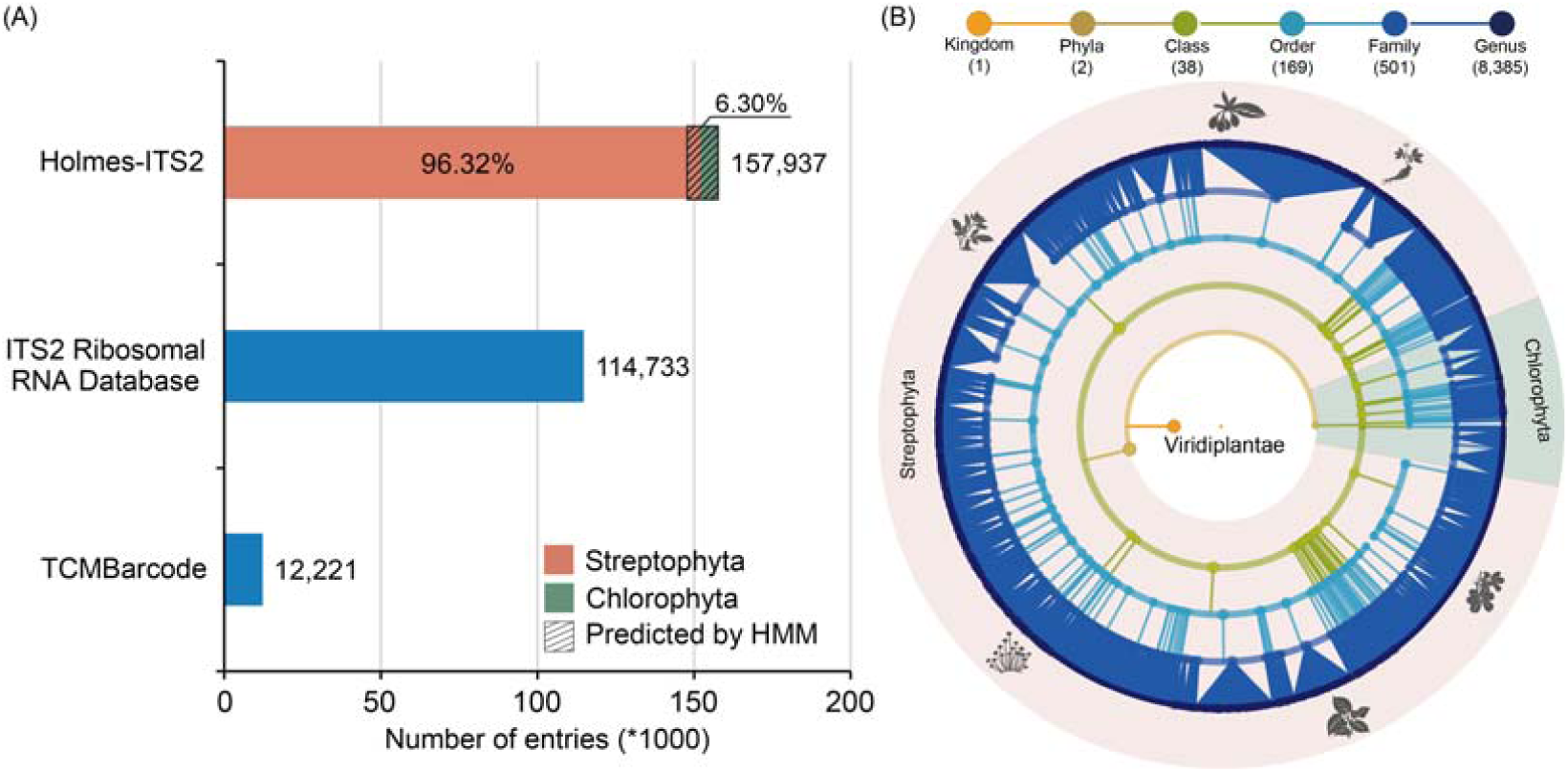
Comparison of data coverage of three databases and taxonomy hierarchical structure of the Holmes-ITS2. **(A)** The number of entries (plant DNA-based marker sequences) contained in each database (blue) and data component of Holmes-ITS2 (red and green). **(B)** From inside to outside of the taxonomy hierarchical tree, each node representing a taxon with a different level of taxonomy such as kingdom, phylum, class, etc. and each interconnect representing an affiliation of the outer node to the inner node.

### Case study in TCM research

For 27 LDW samples(21) we have tested in this study, there were 30,579 ITS2 sequences passing the quality control, with an average of 1,019 sequences of each sample. With sequences passing the quality control and sorted by tag sequences, the biological component identification of each sample based on ITS2 was performed in accordance with the standard operating procedure of Holmes-ITS2 with BLAST for better identification accuracy. The results were summarized in **Table 2**, and it should be noticed that the abundance of a species within a sample depended on both the amount of biological ingredients in that sample and the quality and concentration of DNA during the experiment.

**Table 2.**
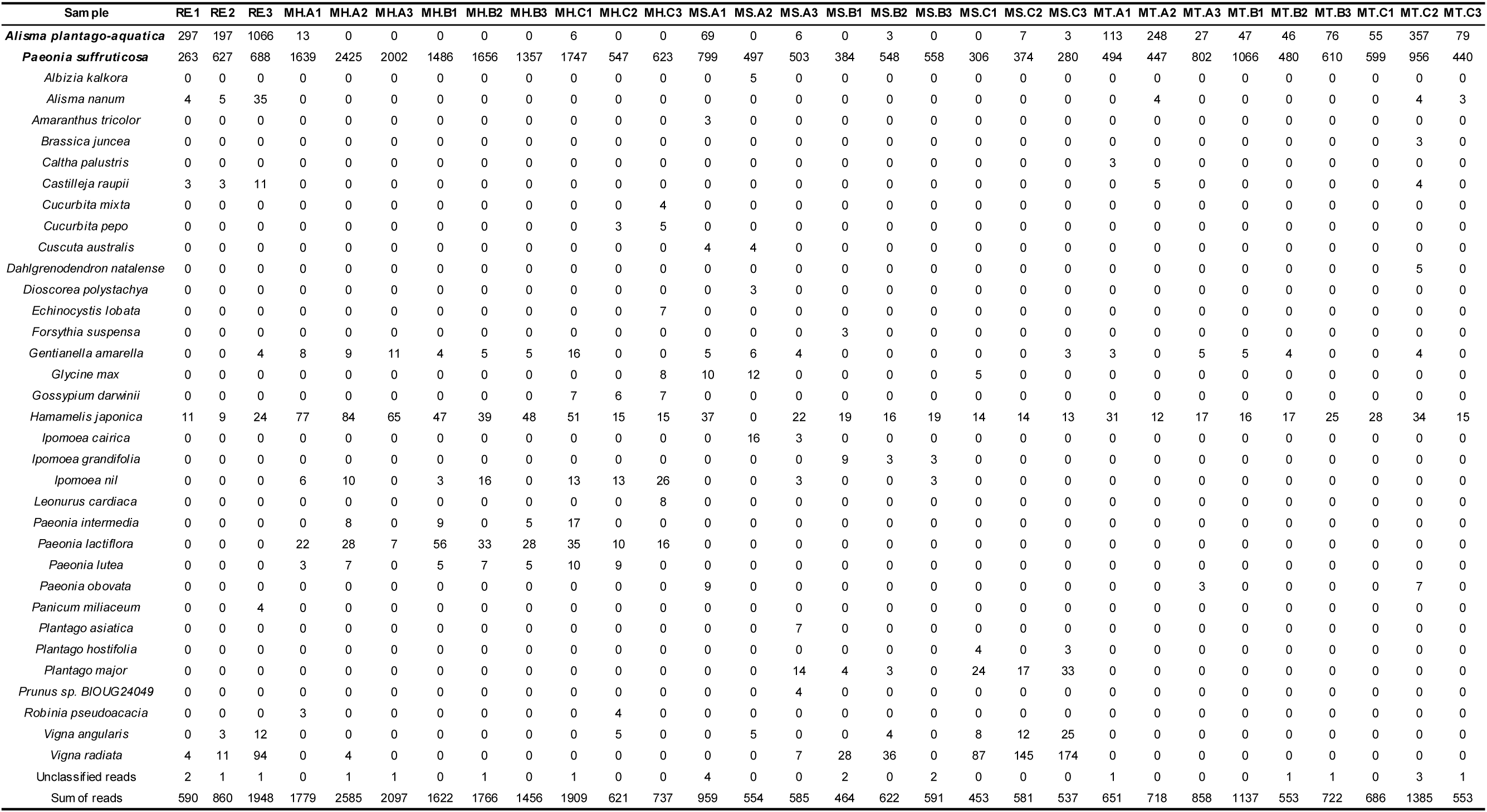
The biological components identified in selected LDW samples based on ITS2 through Holmes-ITS2

In general, compared with the previous identification results based on the raw ITS2 sequences from NCBI as local BLAST database, Holmes-ITS2 database ensured a higher recognition success rate and resolution. Compared to the previous study, in which all unknown sequences could be classified at the genus’s level with partial sequences at the species level, the results of species identification provided by Holmes-ITS2 were almost completely pinned to the species level (The non-identifiable sequences accounted for only 0.075% of all sequences, and half of the samples obtained complete species-level identification results). For example, for sample RE1, the related species *Alisma nanum* of the prescription species *Alisma plantago-aquatica* was detected, which was also detected in RE1’s biologically repeated samples RE2 and RE3, demonstrating its true presence. Similarly, *Castilleja raupii* and *Hamamelis japonica* were also detected in addition to the previously identified species, and other samples were similar. This resulted from the high-quality reference sequences obtained from raw NCBI data processed in the database and the preprocessing of clean ITS2 sequence’s extraction of unknown sequences to be classified through HMM models. In summary, Holmes-ITS2-based biological ingredient identification of TCM preparation could achieve higher identification and success rate (The overall identification results could be accurate to species level with higher resolution of related species) than the direct use of BLAST to search for the original NCBI genbank database. This contributed to the identification of adulterant species and impurities in TCM preparation, demonstrating the availability of Holmes-ITS2 database in actually researches, and analysis of biological ingredients based on ITS2 sequences.

### Web service evaluations

The web service of Holmes-ITS2 was accessible at http://its2.tcm.microbioinformatics.org. The core function of the web service was organism identification and taxonomy classification of plant species based upon the ITS2 sequence. Namely, plant-related ITS2 sequencing data submitted and extracted by the HMM model, were retrieved from the background database through the optional search engine and then species annotation was presented for each submitted sequence. Each batch of data submitted was treated as a sample, and the service also provided statistical information of abundance of the species detected for the sample containing bulk data. Detailed species annotation results were displayed in a tabular form, supplemented by the statistical charts. The database also provided retrieval and browsing of the relevant original sequences and multiple query entries to ensure easy access to query, retrieve and filter information.

The main interface of the web service was shown as **Figure 7** with main browse and search functional pages. In browse page, all species sequences could be browsed, and their corresponding species’ phylonenetic positions and detailed descriptions were shown and could be linked-out to their Wikipedia pages (**Figure 7 (A)**). In search page, after submission of multiple sequences through the “Enter Query Sequence” window, selection of appropriate search engine by switching the label and corresponding DNA barcode dataset as background dataset, the species identification of the target sequences could be initiated (**Figure 7 (B)**).

**Figure 7.**
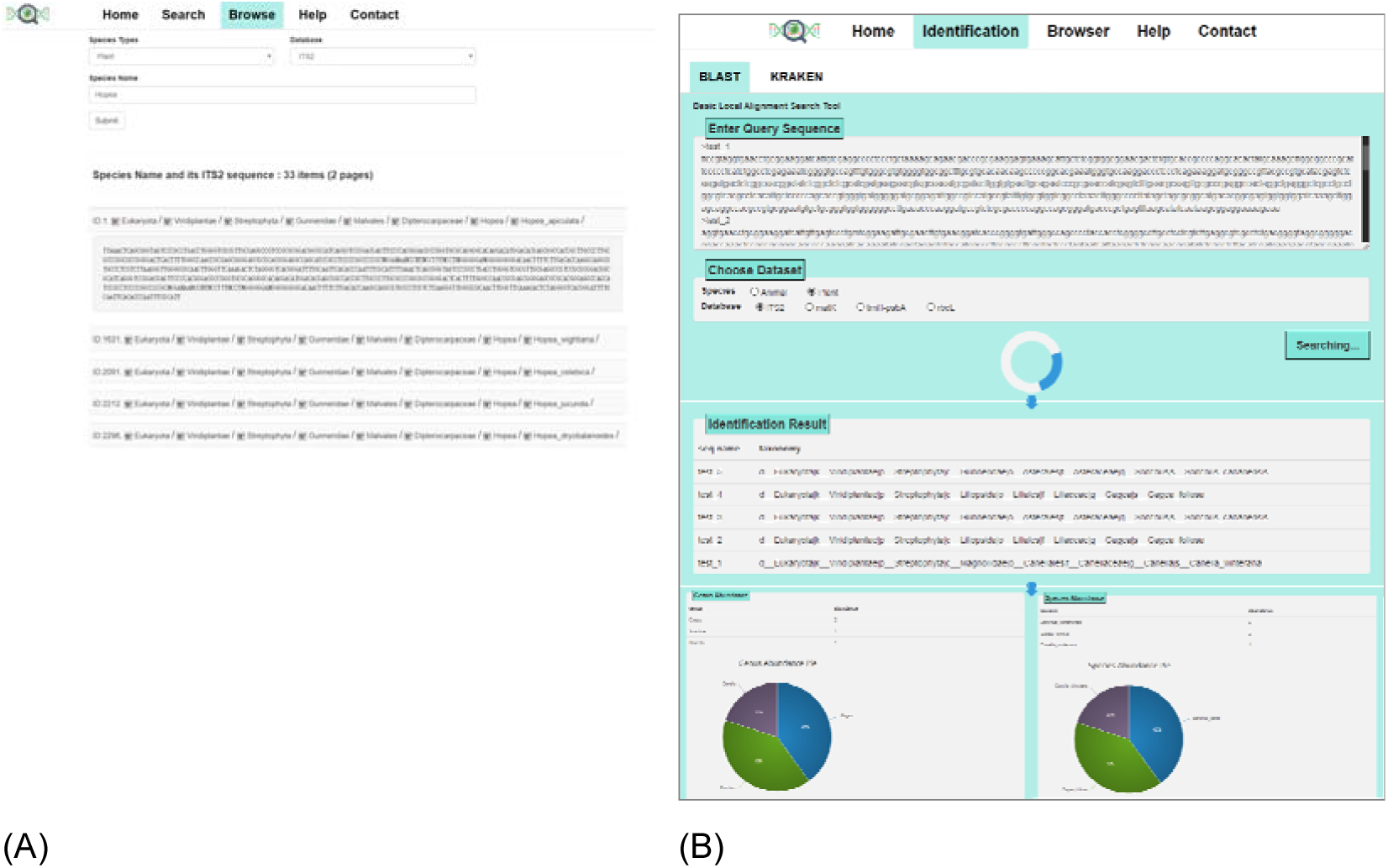
The interface of the Holmes-ITS2 web services. (A) All species sequences could be browsed, and their corresponding species’ phylonenetic positions and detailed descriptions were shown with link-outs. (B) BLAST and Kraken search algorithms were available and multiple DNA-based markers were optional.

Species identification results included the species information for each sequence displayed in a tabular form, and the species with top ten abundance of the sample at the taxonomy level of species and genus displayed in tables and pie charts, and showed the abundance composition of the entire sample (**Figure 7 (A)**).

### Evaluation of data updates mechanism

In the last ten years, the number of ITS2 sequencing data rose significantly and steadily compared with the past and showed a rising trend for future researches (**Figure 8**). Thus, it was essential to continue tracking the new data released. With the data update mechanism of Holmes-ITS2, the data when we have performed our latest update was in May 2017. With the entries meeting the standards set and refined through the clustering approach, there were 17,436 sequences newly added, which belongs to 7,139 new species. With this update performed, ITS2 entries in Holmes-ITS2 would keep up to date to ensure the maximum data coverage.

**Figure 8.**
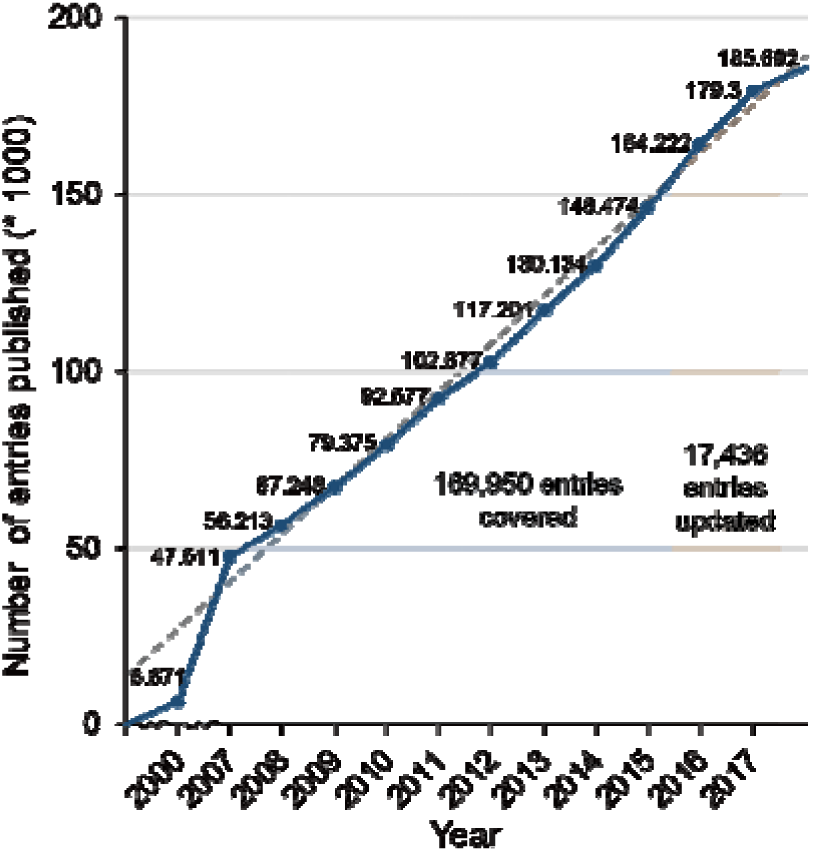
Increase of the number of ITS2 entries in Holmes-ITS2, extracted from NCBI on yearly basis. Data published (searched with key word ‘ITS2’ in the NCBI nucleotide database) in the last ten years showed a significant rise compared with the past and a steady upward trend in the future, and 17,436 entries newly released until May 2017 were covered.

## DISCUSSIONS AND CONCLUSION

Based on the ITS2 sequences of plants, this study constructed a plant DNA-based marker database (Holmes-ITS2), aiming at organismal identification and taxonomic classification of plant species or plant mixed systems including but not limited to Chinese herbal medicine or TCM preparations in actual researches, which is significance in evaluation of the efficacy and safety of this kind medicine for human use(23). Compared with the existing databases, this work improved the classification performance in terms of accuracy, efficiency and data coverage, especially for large amounts of data produced by high-throughput sequencing technology, providing accurate and high efficient analysis towards plant species identification.

To be specific, with the help of ITS2 HMM model constructed, it would be easier for the extraction of ITS2 regions of sequences obtained from experiments that aiming at amplification of the whole ITS fragment. Source of variation within ITS region was mainly depended on the ITS2, hence accuracy of taxonomy classification based on ITS2 was improved. In addition, the comprehensive pretreatment of raw entries in aspects of both sequences and taxonomy annotations, including strict ITS2 region extraction, etc. made it reliable reference as in the Holmes-ITS2. In consideration of time efficiency of taxonomy classification in actual use, multiple search engines were assessed, and the best-performing algorithms were adopted for the relatively high accuracy and efficiency. In terms of classification accuracy of ITS2 sequences, which was the main concern, Holmes-ITS2 showed certain advantages over the two existing databases, including TCMBarcode and ITS2 Ribosome Database. In genus’s level, the accuracy of Holmes-ITS2 was over 17% higher than the two existing databases in average, while undermost 10% advantage in species-level, which ensured the reliability of classification results based on Holmes-ITS2. As data generated by next-generation sequencing technique was booming, which raised a high demand of efficiency of data analysis strategies, Holmes-ITS2 overcame the short board that existing service’s lacking of handling a large batch of data supported by the high-performance computing platform. As the test result, classification time cost could maintain a linear increase trend, which made the time cost predictable. Overall, Holmes-ITS2 achieved the design goals, including the improvements in aspects of both accuracy and efficiency basically.

According to the results of classification accuracy of all three databases to be compared, it was found that the resolving power of taxonomy classification based on ITS2 as nucleotide DNA-based marker was relatively high to genus level in general (over 94% of Holmes-ITS2). In pursuit of the species-level precision, the performance of ITS2 as DNA-based marker didn’t give complete satisfaction. It was inferred that it may be caused by the fact that the average length of ITS2 sequence was short (about 200 base pairs) so that it couldn’t provide enough variation in evolution to differentiate partial plant species. The cluster analysis also indicated that the relatively high similarity between certain species with far phylogenetic relationship. Therefore, as a supplement, other nucleotide DNA-based markers of plants including matK(24), rbcL(25), psbA-trnH(26) was planned to be brought into Holmes-ITS2 database as supplement (there existing such problems, including non-homology (matK) and heterogeneity that prevent the creation of a universal PCR toolkit (rbcL)). Besides, in consideration of the actual experiments, combinatorial markers would complement the shortage of each marker, which lead the discovery of more existing species. Furthermore, to keep up on the latest researches and data published to ensure the maximum species coverage, the data update mechanism would be performed at least once per around half year, and the entire database and web server would be maintained actively for the maximize real-time availability.

Our goal for development of Holmes-ITS2 database has been to develop a high accurate, high efficient and comprehensive species coverage organism identification and taxonomy classification system with a user friendly and highly available platform (web service). As the diversity of data and methods, the next step in development of Holmes-ITS2 focuses on two aspects: (1) collection and processing of DNA-based marker data, including ITS2 and more nucleotide DNA-based markers as supplement and more accurate and efficient matching search engines, and (2) the integration of related downstream preliminary statistical analysis tools. This effort will advance the utility of Holmes-ITS2 and increase its value as a taxonomy identification platform of plant or plant mixed system, including herbal medicine or TCM preparations.

## ACKNOWLEDGEMENT

This work is partially supported by National Science Foundation of China grant 31671374, Ministry of Science and Technology’s high-tech (863) grant 2014AA021502, and Sino-German Research Center grant GZ878.

## COMPETING FINANCIAL INTERESTS

The authors declare no competing financial interests.

